# Designed hybrids facilitate efficient generation of high-resolution linkage maps

**DOI:** 10.1101/243451

**Authors:** Kazutoshi Yoshitake, Yoji Igarashi, Misaki Mizukoshi, Shigeharu Kinoshita, Susumu Mitsuyama, Yutaka Suzuki, Kazuyoshi Saito, Shugo Watabe, Shuichi Asakawa

**Affiliations:** Laboratory of Aquatic Molecular Biology and Biotechnology, Graduate School of Agricultural and Life Sciences, The University of Tokyo, 1-1-1 Yayoi, Bunkyo, Tokyo 113-8657, Japan.; Department of Medical Genome Sciences, Graduate School of Frontier Sciences, The University of Tokyo, 5-1-5 Kashiwanoha, Kashiwa, Chiba 277-8561, Japan.; Akita Prefectural Institute of Fisheries, Oga, Akita 010-0531, Japan.; School of Marine Biosciences, Kitasato University, Sagamihara, Kanagawa 252-0373, Japan

**Keywords:** linkage map, scaffold extender, hybrid organisms, whole genome SNP typing, *Takifugu rubripes*, *Takifugu stictonotus*

## Abstract

In sequencing eukaryotic genomes, linkage maps are indispensable for building scaffolds with which to assemble and/or to validate chromosomes. However, current approaches to construct linkage maps are limited by marker density and cost-effectiveness, especially for wild organisms. We have now devised a new strategy based on artificially generated hybrid organisms to acquire ultra high-density genomic markers at lower cost and build highly accurate linkage maps. Using this method, linkage maps and draft sequences for two species of pufferfish were obtained simultaneously. We anticipate that the method will accelerate genomic analysis of sexually reproducing organisms.

---

New-generation technologies now enable whole-genome sequencing of any organism, although assembling long, high-quality genomes from short reads generated by these technologies remains challenging. Accordingly, methods such as Irys genome mapping (BioNano Genomics Inc.), Hi-C, Chicago library (Dovetail) and linked reads (10× Genomics) have been developed to generate longer scaffolds and improve assembly(Burton et al. 2013; Putnam et al. 2016; Bickhart et al. 2017; Weisenfeld et al. 2017). Nevertheless, genetic linkage analysis would likely remain essential to reconstruct most eukaryotic genomes, even if sequencing methods continue to advance. Indeed, linkage maps are used not only to assemble contigs/scaffolds, but also to compare and evaluate the assembled sequences.

Construction of conventional linkage maps based on microsatellite markers generally is costly and time-consuming, and the maps are of low resolution and genomic coverage due to the limited number of markers, usually several hundreds to several thousands(Kai et al. 2011). Several new methods to construct linkage maps and assemble contigs/scaffolds have also been developed, including SNP array(Bodenes et al. 2016), RAD-seq(Arora et al. 2017), and genotyping-by-sequencing(Nunes et al. 2017). However, SNP arrays require SNP data and hundreds of arrays prepared in advance, and therefore may not be suitable for *de novo* genome analysis. In contrast tens of thousands of SNP markers are easily obtained by RAD-seq or genotyping-by-sequencing, although the distribution of such markers strongly depends on the restriction enzymes used. In addition, markers obtained from these methods are usually present at one per tens of kb; thus, they lie outside the length of most contigs and small scaffolds, and cannot be located on the genome.

Determining the two haplotypes in each individual is the main challenge in linkage analysis (Fig. 1A). Indeed, typing of polymorphic sites by genome sequencing generally requires coverage higher than 10 fold, on average, to distinguish between hetero- and homozygosity. As several hundred individuals are usually genotyped, a large volume of tasks would be required to achieve the required coverage for each individual. Such large tasks may thus prevent the use of whole-genome sequencing for this purpose, even if throughput from sequencing technologies significantly expands.

**Figure 1.**
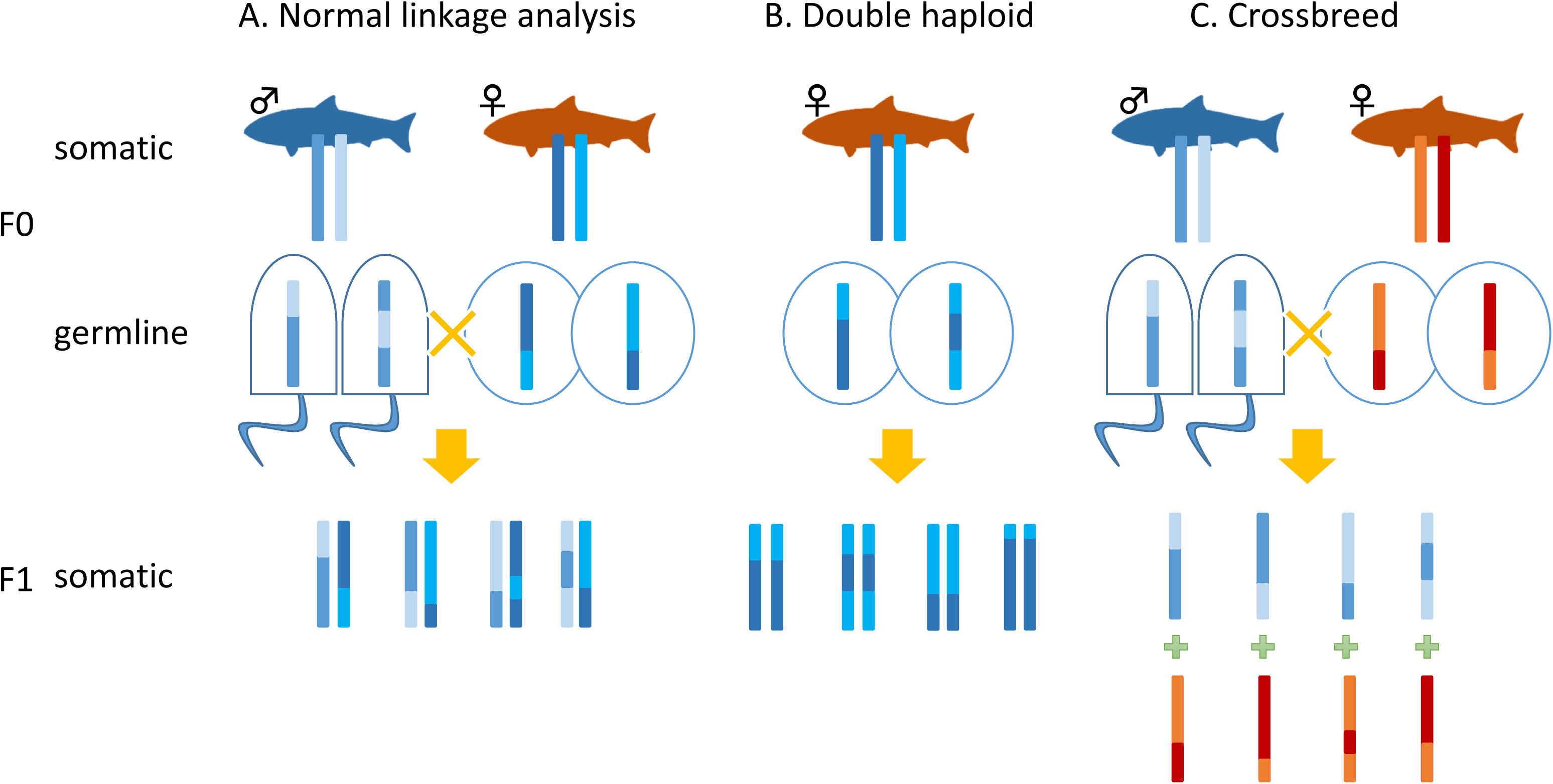
Comparison of linkage analysis methods. The types of individuals necessary for linkage analysis to extend genomic sequences are shown for each method. **(A)** Crosses and chromosomes for linkage analysis in normal wild populations. **(B)** Chromosomes of double haploid individuals. **(C)** Chromosomes of hybrid individuals.

However, a single read of each polymorphic site is sufficient to determine the haplotype in haploid individuals, such as those generated by gynogenesis or androgenesis in some organisms (Fig. 1B), and each individual can thus be haplotyped with substantially fewer reads. The haplotype inherited from the diploid parent can then be inferred. By extension, haplotyping is also very simple if each read can be clearly attributed to the mother or father (Fig. 2A). For example, when we find a read inherited from the mother, we need only determine either of the two maternal haplotypes, for which the required coverage is only one (Fig. 1C). By this strategy, independent linkage maps can be generated for both the mother and father (Fig. 1C, Fig. 2C). However, maternal and paternal genomes are typically almost the same and are usually indistinguishable in the offspring.

**Figure 2.**
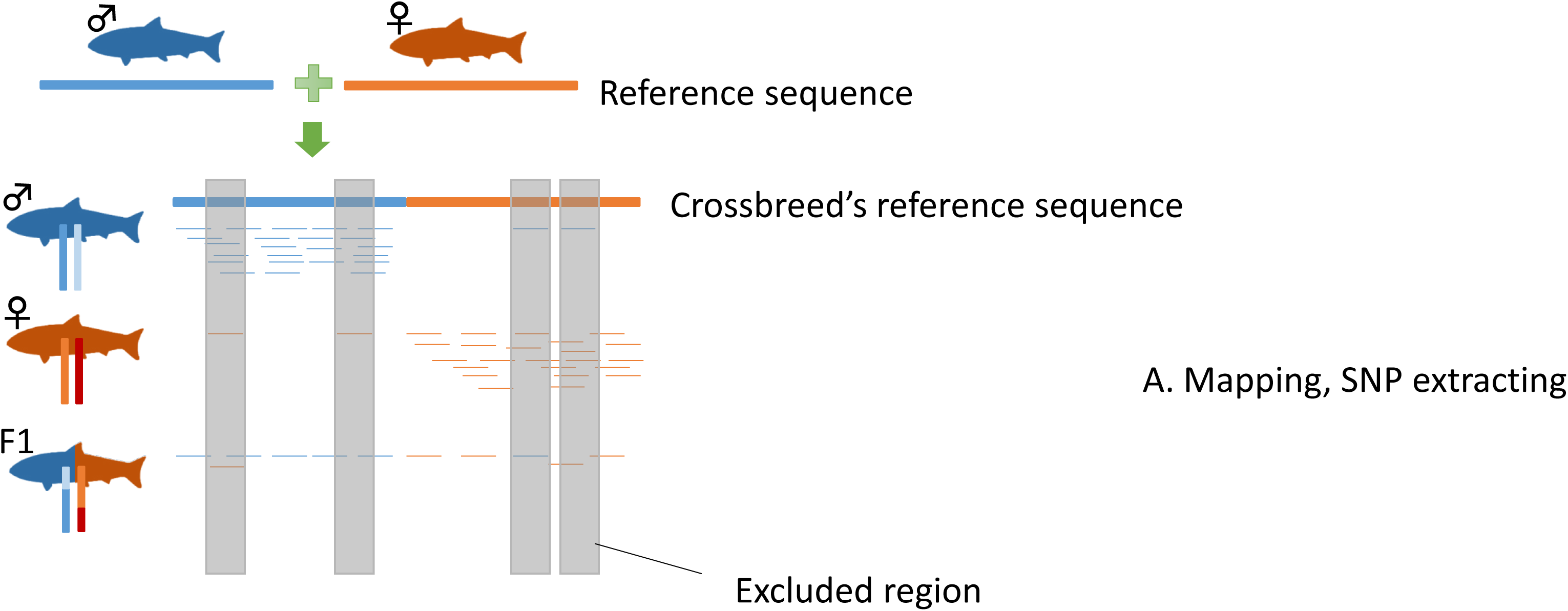
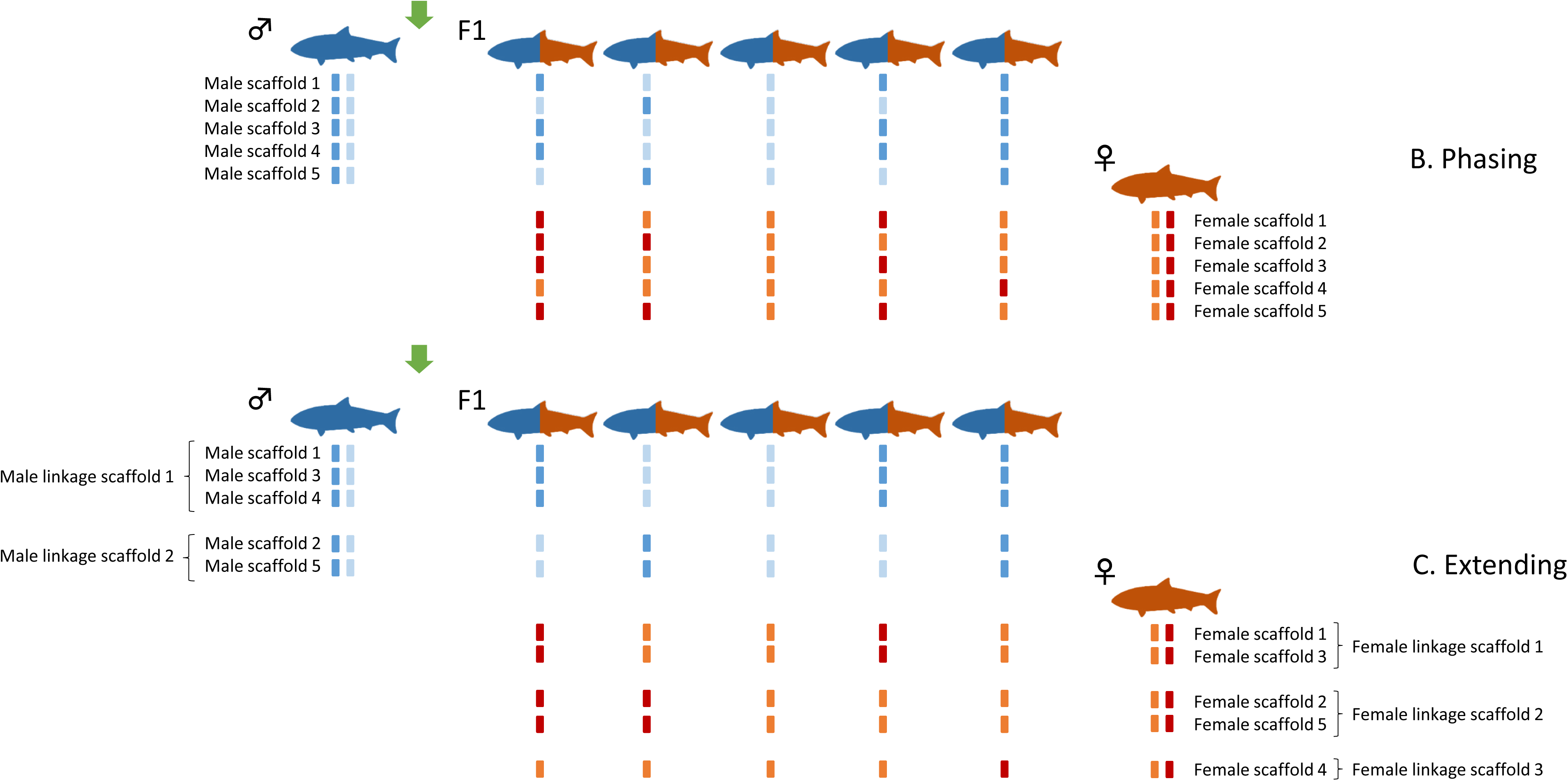
SELDLA. **(A)** Sequence data are mapped to a reference sequence obtained by combining parental genomes, and SNPs in specific regions are extracted. **(B)** To analyze the genome of a parent, we focus only on the reference sequence of that parent, determine which of its two chromosomes was passed to the hybrid, and phase. This is repeated from the beginning of the scaffold. **(C)** We arrange scaffolds with similar phase patterns on the chromosome. If phase patterns are different at the beginning and end of the scaffold, it is also possible to determine the orientation on the chromosome.

To test this approach, we generated hybrid offspring from two infertile species (Fig. 1C), one of which is torafugu, or tiger pufferfish *(Takifugu rubripes*), and the other is gomafugu *(Takifugu stictonotus). T. rubripes* is suitable as a reference, as a draft genome and linkage map are already available. In contrast, only the mitochondrial genome is known for *T. stictonotus.* As the analysis is based on whole genomes, almost all SNPs are suitable as polymorphic markers, the density of which are typically higher in comparison to all others. Accordingly, the resolution of the resulting linkage map would be limited in theory not by the number of markers, but by the number of crossover sites.

The linkage maps thus obtained simultaneously for both species were high-resolution and high-coverage, especially in comparison to the latest draft sequence for *T. rubripes* (FUGU5). We also constructed the first linkage map and draft genome sequence for *T. stictonotus,* consisting of 22 chromosomes. We believe this is the first instance in which a draft sequence of the correct number of chromosomes was obtained in a single analysis. Our strategy should be applicable to most sexually reproducing organisms, and thus should facilitate genome analysis of not only oviparous organisms, but also viviparous animals and plants.

## RESULTS

### Sequencing of samples

We sequenced 192 hybrid fry, and obtained suitable data from 188. Sequence data were also obtained from the father, as well as from a mother-equivalent following loss of the mother. Sequence statistics are listed in Table 1. Assuming a genome size of 391 Mb for both species, coverage was 60.7 and 83.5 fold for *T. rubripes* and *T. stictonotus,* respectively. Because the size of the genome is doubled (782 Mb) in hybrids, coverage for each was 1.8 fold on average.

**Table 1.**
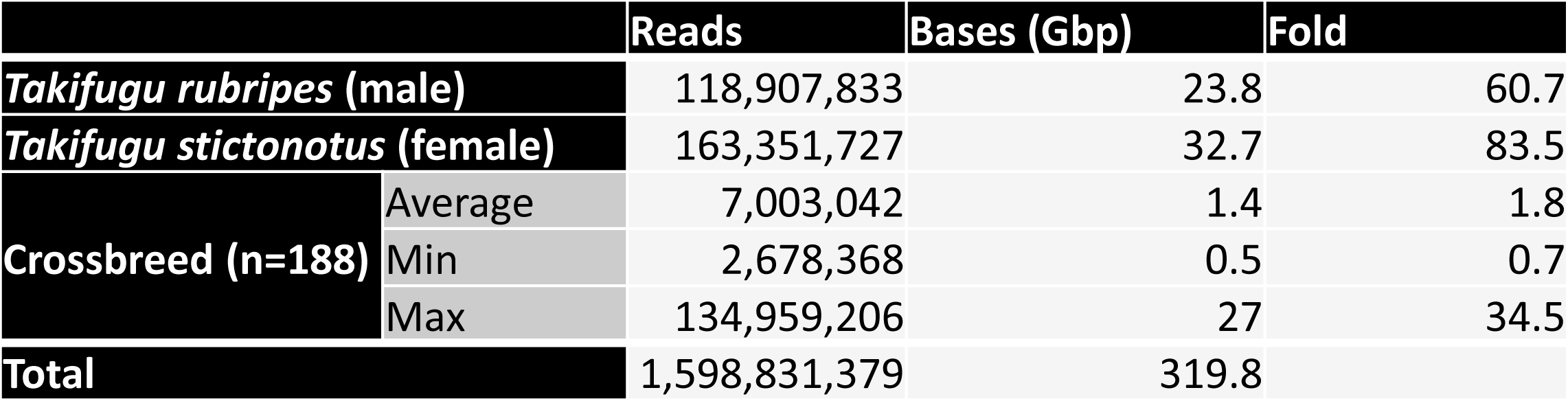
Sequencing statistics for the male parent *(T. rubripes),* a female *T. stictonotus,* and hybrid fry. To calculate depth, a genome size of 391 Mb was used for *T. rubripes* and *T. stictonotus,* as previously reported. The genome size was doubled to 782 Mb for hybrid fry.

|  |  | Reads | Bases (Gbp) | Fold |
| --- | --- | --- | --- | --- |
| ***Takifugu rubripes* (male)** |  | 118,907,833 | 23.8 | 60.7 |
| ***Takifugu stictonotus* (female)** |  | 163,351,727 | 32.7 | 83.5 |
| **Crossbreed (n=188)** | Average | 7,003,042 | 1.4 | 1.8 |
|  | Min | 2,678,368 | 0.5 | 0.7 |
|  | Max | 134,959,206 | 27 | 34.5 |
| **Total** |  | 1,598,831,379 | 319.8 |  |

### Reference genome assemblies

The N50 for the *de novo T. rubripes* genome that we obtained (de-novo-contigs-Tr) was 13.0 kb (Table 2A), which is 66 times shorter than that of the previous version of the *T. rubripes* genome (FUGU4), for which the N50 was 858 kb. Therefore, FUGU4 was used as a reference in further analysis. For *T. stictonotus,* the genome that we obtained (de-novo-contigs-Ts), with N50 of 15.9 kb, was used as a reference because no other genome sequence is available.

**Table 2.**
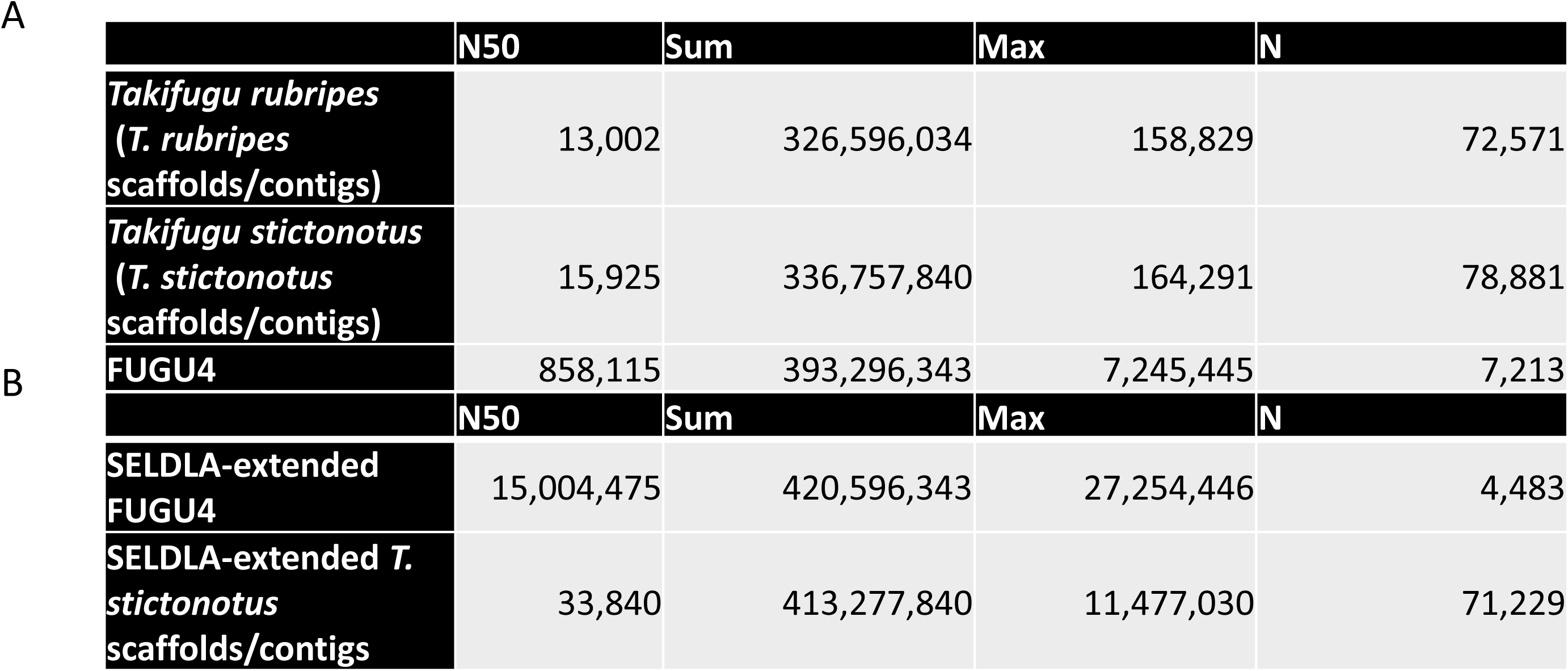
Assembly and scaffolding. **(A)** Assembly statistics for *de novo T. rubripes* and *T. stictonotus* genomes, as well as for FUGU4 and FUGU5, which are published *T. rubripes* genomes. N50, the shortest sequence length at 50 % of the genome; sum, total scaffold length; max, maximum scaffold length; n, number of scaffolds. **(B)** Assembly statistics for SELDLA-extended FUGU4 and SELDLA-extended *T. stictonotus* genome.Table 1 Outline of sequence data Table 2 The result of assemble and scaffolding

### Extraction of SNP markers from very low-coverage data

We developed Scaffold Extender with Low Depth Linkage Analysis (SELDLA), an analysis pipeline (Fig. 2) with which we attempted to extend FUGU4 scaffolds using new low-coverage sequence data from hybrid fry. We then assembled the extended FUGU4 contigs/scaffolds according to the new SELDLA-derived linkage map, and compared the assembly with FUGU5, the current version of the *T. rubripes* genome, which was constructed by ordering FUGU4 contigs/scaffolds according to a linkage map of 1,222 microsatellites. We regarded the FUGU4 plus de-novo-contigs-Ts as a haploid reference sequence for a hybrid individual. FUGU4 and the de-novo-contigs-Ts genome were 98.1 % homologous over aligned sequences throughout the genome. We then mapped each read of the hybrid individual to the FUGU4 plus de-novo-contigs-Ts genome using BWA mem (Li and Durbin 2010), and extracted those in which a continuous stretch of 90 bp or more (out of 100 bp) was mapped exclusively and uniquely. The average mapping rate after this process was 82.8 %. Subsequently, SNPs were called in GATK UnifiedGenotyper(McKenna et al. 2010). We note that SNPs were called even if present in only one read, which is generally discarded as a sequence error in conventional analysis.

### Extraction of SNP markers form *T. rubripes*

Linkage analysis was performed separately for *T. rubripes* and *T. stictonotus.* In the former, 5,140,280 SNPs were initially extracted. However, a site was regarded as highly similar between the two species and thus excluded from linkage analysis, if even a single *T. stictonotus* read was mapped to *T. rubripes* (Fig. 2A). As a result, only 1,293,394 SNPs remained for further processing. To minimize potential errors, reads mapped with a depth more than 4 times the average depth were then removed. Furthermore, we extracted SNPs confirmed to be heterozygous in the paternal genome, as every SNP found in paternal sequences in hybrid fry should also be found in the diploid genome of the father. After these steps, 606,110 markers were available for analysis. SNPs found in less than 10 % of samples were excluded, along with those for which minor alleles were found in less than 30 % of samples (major alleles found in more than 70% of samples). SNPs heterozygous in more than 20 % of individuals were suspected to be multi-copy genes, and were also filtered out. Finally, SNPs that appeared to be a crossover breakpoint in a scaffold/contig of more than 10 % of individuals were regarded as noise and were excluded. Ultimately, 442,723 markers were used in linkage analysis, for an average density of one per 903 bp.

### Elongation of FUGU4 scaffolds by SELDLA

Because algorithms to order and orient scaffolds are known to be NP-hard(Pop et al. 2004; Tang et al. 2015), a normal heuristic approach was applied. Briefly, each hybrid fry was phased from the beginning of a scaffold, and phase changes in the middle of the scaffold were listed as break points. After end-to-end phase patterns were collected from 188 individuals, scaffolds were sorted in descending order of length. Scaffolds with the most similar phase patterns to FUGU4 were then connected to FUGU4 scaffolds at both ends. This process was repeated for the resulting connected scaffolds, until no test scaffold was found for which the similarity to a target scaffold was above the threshold value. If the phases at both ends of the newly connected scaffold were the same, the orientation of the scaffold could not be determined, and was either filled with Ns (any of A, C, G, T bases) along the length of the unoriented scaffold, or was used without determining the orientation. Both approaches were used in this study.

Using 442,723 SNP markers, we phased all 188 *T. rubripes* genomes, as well as 4,513 of 7,213 scaffolds in FUGU4, such that phase was obtained for 95.7 % of all nucleotides in the genome. Orientation was determined for 834 scaffolds representing 74.8 % of nucleotides in the genome.

### Comparison of FUGU5 and SELDLA-extended FUGU4

As listed in Table 2B, extension by SELDLA increased the N50 in FUGU4 by 17 fold, such that it was now longer than that of FUGU5, which was generated by extending FUGU4 using conventional linkage analysis. A Circos plot (Fig. 3) confirmed that the genomic organization in SELDLA-extended FUGU4 is consistent with that of FUGU5 over 22 chromosomes, except at one site. Dot plots comparing all scaffolds between FUGU5 and SELDLA-extended FUGU4 are shown in Fig. 4 and Supplementary Fig. 1, and indicate that alignment was mostly successful throughout the entire genome. In addition, we found that many small scaffolds/contigs unmapped in the former were aligned at both ends to chromosomes in the latter (Fig. 4 and Supplementary Figs. 1), as clearly shown in a zoomed-in view of chromosome 1, a representative example. Indeed, several small scaffolds/contigs were incorporated and ordered into chromosome 1 in SELDLA-extended FUGU4, but not in FUGU5. This result implies that microsatellite markers near telomeric regions were inadequate in FUGU5, and highlights the large difference in marker density between microsatellites in FUGU5 and non-selected SNPs in this study. Several genomic inversions and translocations were also observed, suggesting structural polymorphisms between *T. rubripes* individuals. We note that these may cause erroneous ordering in SELDLA-extended FUGU4, FUGU5, or both. Finally, comparison of physical and linkage distance (Supplementary Fig. 2) indicated that linkage distance was tend to be longer around telomeres, confirming that our result was consistent with previous study(Kai et al. 2011).

**Figure 3.**
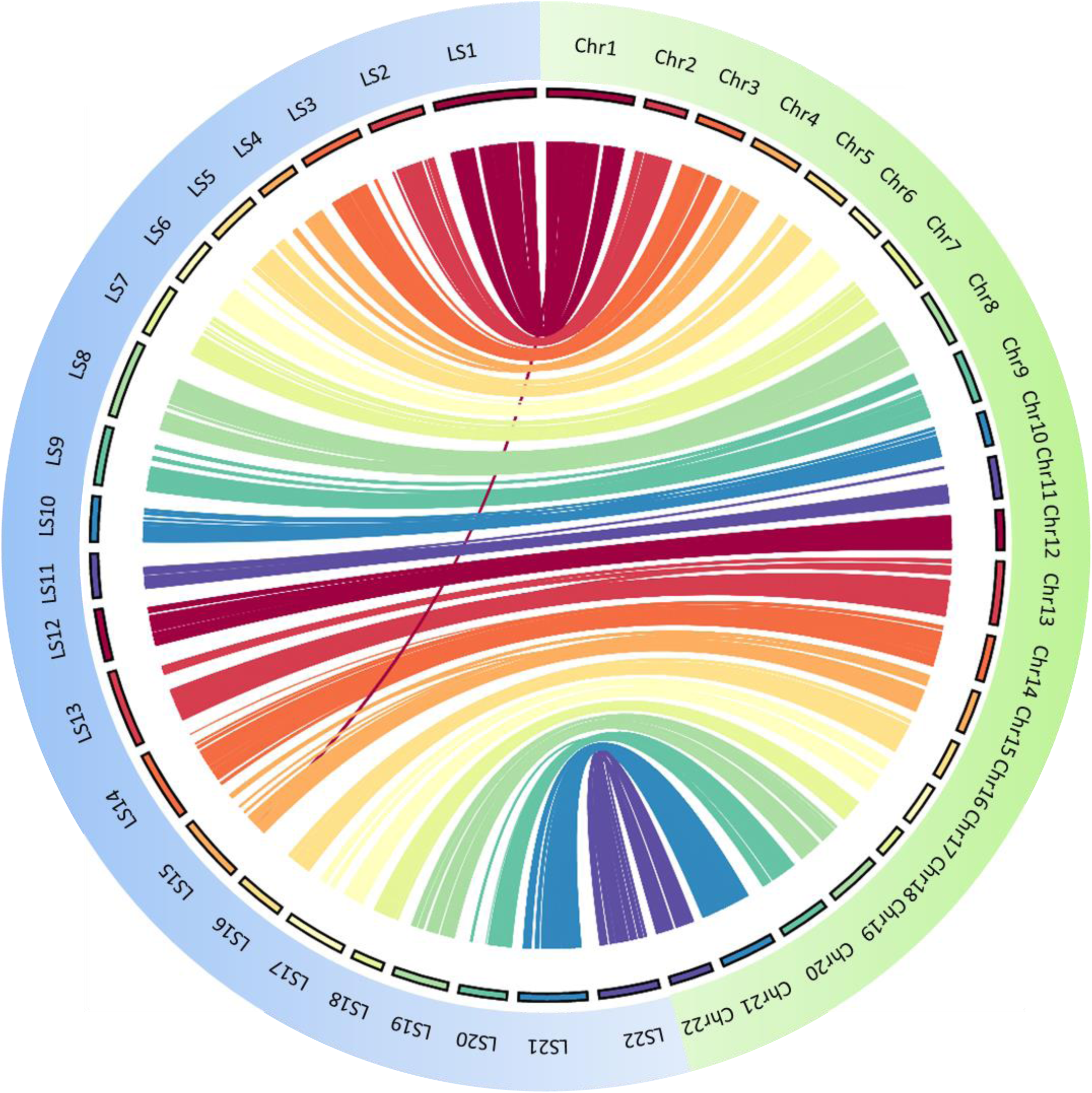
Circos comparison of 22 *T. rubripes* chromosomes between SELDLA-extended FUGU4 and FUGU5. Regions that could be aligned over 1 kb or more are connected by lines. LS: Linkage-group-extended (SELDLA-extended) Scaffold, chr: chromosome of FUGU5.

**Figure 4.**
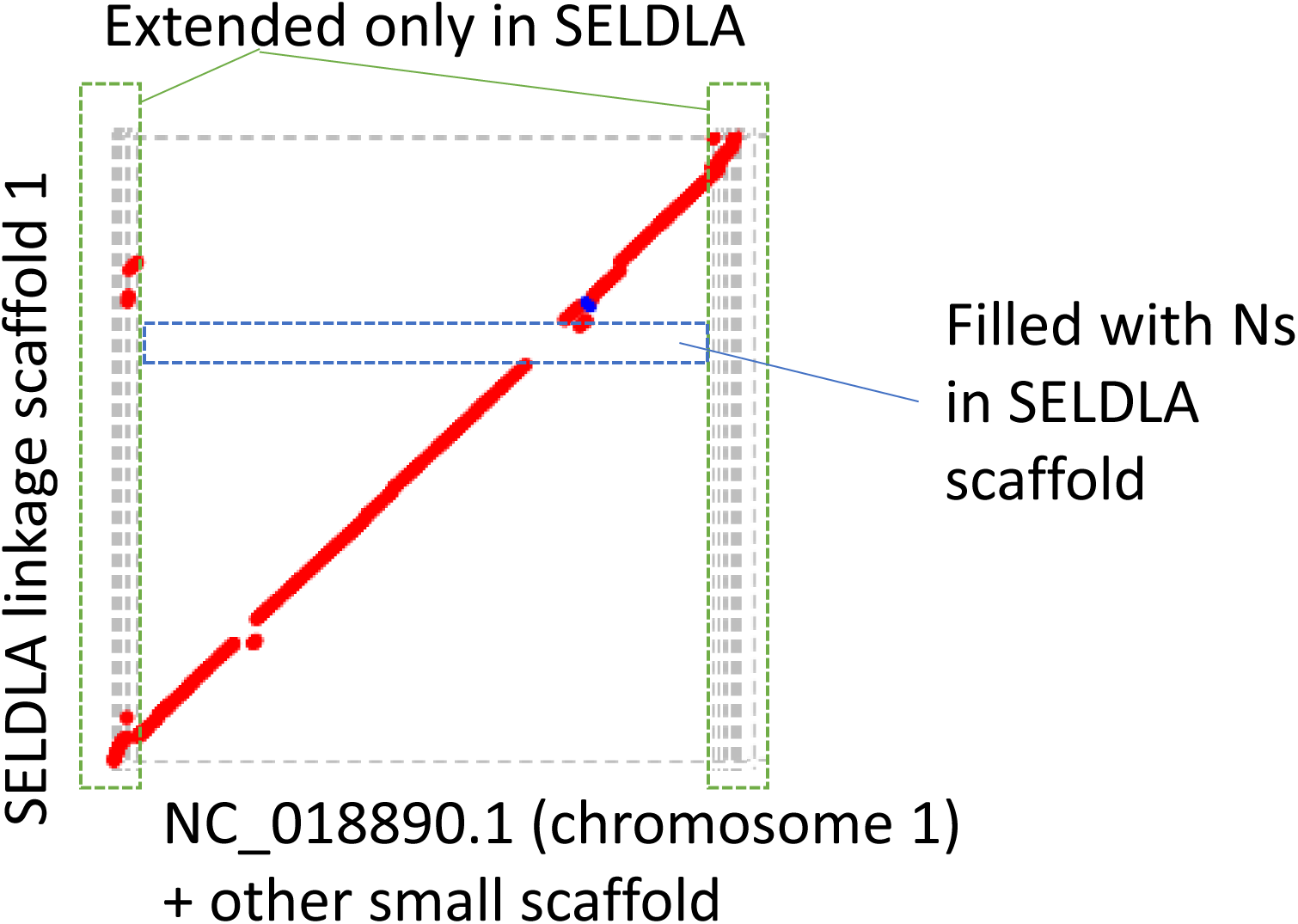
Dot plot homology comparison of chromosome 1 between SELDLA-extended FUGU4 and FUGU 5. Regions that could be aligned over 1 kb or more are plotted, with FUGU5 on the x-axis and SELDLA-extended FUGU4 on the y-axis. Red lines indicate alignment in the forward direction, whereas the blue line indicates alignment in the reverse direction. The plot shows that it was possible to arrange small FUGU5 scaffolds near telomeres according to the SELDLA-extended chromosome.

### Scaffold elongation of the *T. stictonotus* genome by SELDLA

In addition, we also attempted to extend *T. stictonotus* contigs by SELDLA. These contigs were obtained from a *T. stictonotus* female in a single round of paired-end sequencing on Illumina HiSeq2000, followed by assembly in CLC genomics workbench. The N50 for this genome was 15 kb. Although it is generally very difficult to perform linkage analysis using very short and numerous scaffolds/contigs, such analysis was possible for *T. stictonotus* due to the very high density of SNP markers.

By extracting markers according to the same process used for *T. rubripes,* 144,596 SNPs were obtained. Nevertheless, these markers were only 33 % of the amount obtained from *T. rubripes,* perhaps reflecting closer kinship in *T. stictonotus.* Of the 78,881 scaffolds in the draft genome, phases were determined for 10,309, encompassing 22.9 % of all nucleotides. These phased scaffolds were clustered in 22 linkage groups, corresponding to the number of chromosomes/linkage groups in *T. rubripes.* Orientation was also determined for 1,097 scaffolds, covering 7.7 % of all nucleotides. The remaining 9,212 scaffolds/contigs were mapped to linkage groups, but in unknown orientations, mainly because there were no crossover points in the genome corresponding to short scaffolds/contigs from any of 188 individuals. In some cases, this effect might be due to lack of real SNPs. Surprisingly, the maximum scaffold length in the SELDLA-extended genome was 15.9 Mbp, which is 68 % of the maximum scaffold length in FUGU5.

As *T. rubripes* and *T. stictonotus* belong to the same genus, the genome organization of both was expected to be similar. Indeed, a Circos plot of SELDLA-extended *T. stictonotus* and FUGU5 showed not only synteny, but also highly conserved sequence organization (Fig. 5). In addition, whole-genome dot plots (Fig. 6 and Supplementary Fig. 3) showed contiguous alignment over the entire chromosomes, although only 22.9 % of nucleotides were phased. Physical and genetic distances in each SELDLA-extended *T. stictonotus* and FUGU5 are plotted in Supplementary Fig. 4.

**Figure 5.**
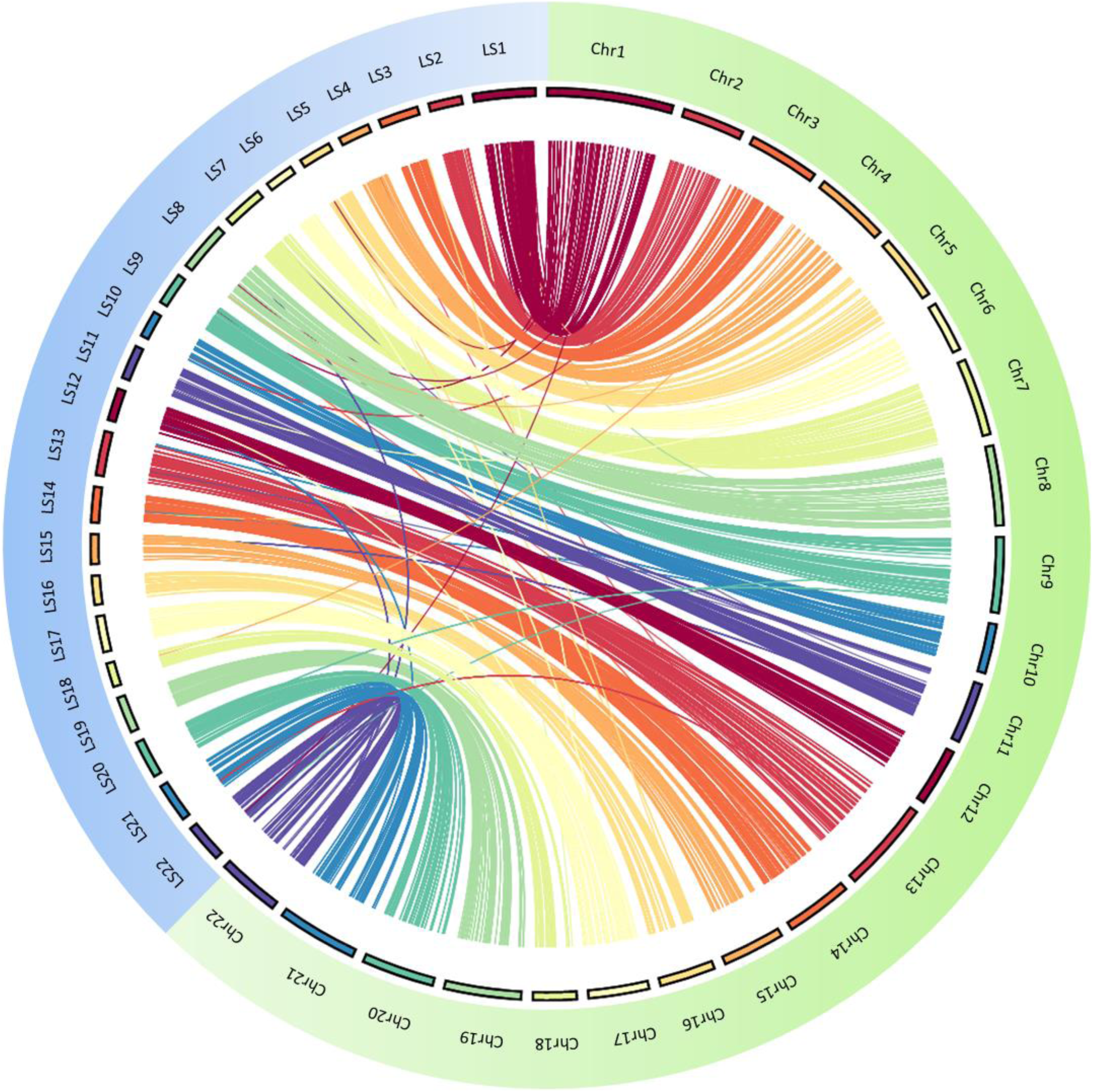
Comparison of FUGU5 and SELDLA-extended *T. stictonotus* genome. Regions that could be aligned over 1 kb or more in each of 22 chromosomes are connected by lines.

**Figure 6.**
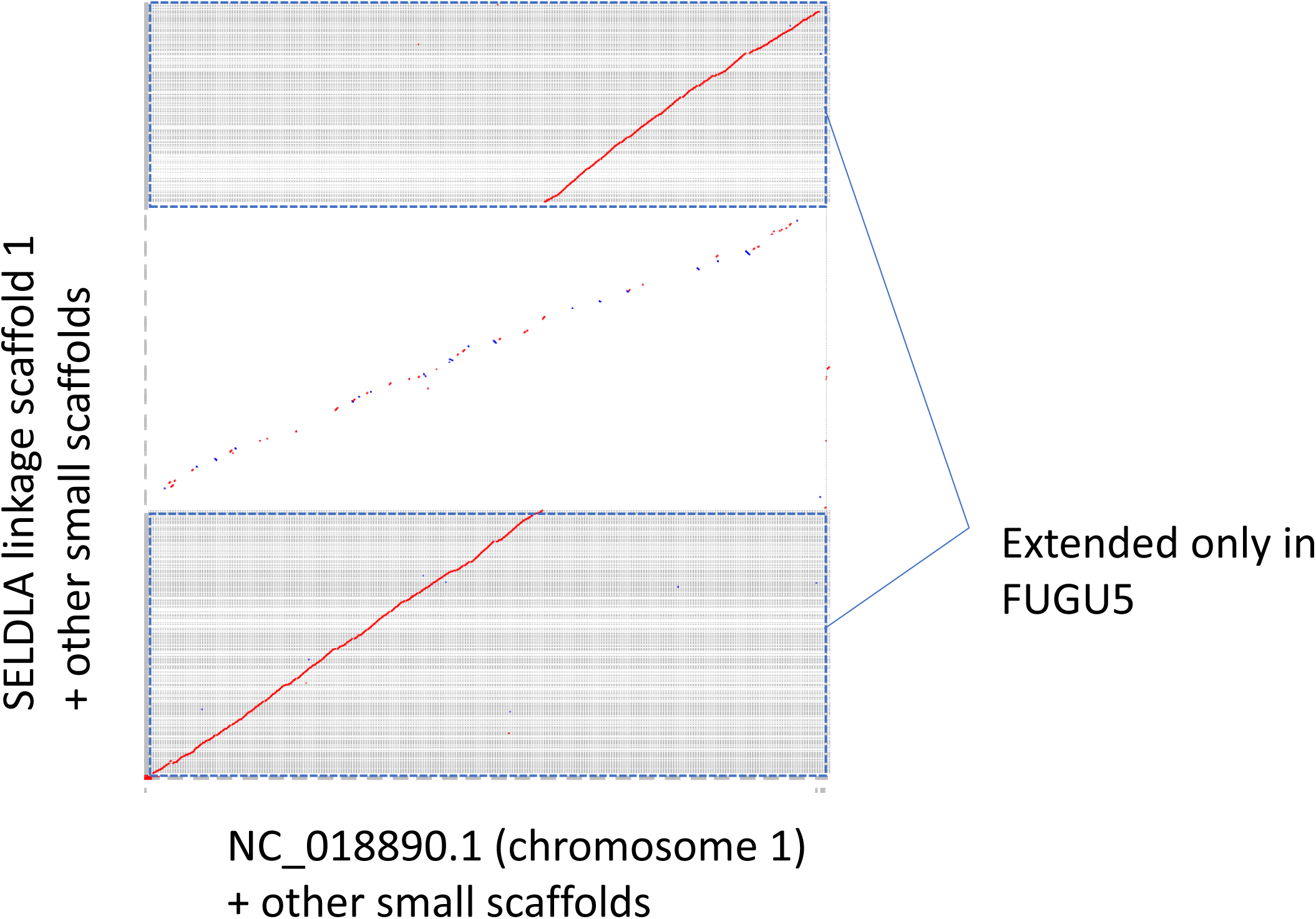
Dot plot homology comparison of chromosome 1 between FUGU 5 and SELDLA-extended *T. stictonotus* genome. Regions that could be aligned over 1 kb or more are plotted, with FUGU5 on the x-axis and SELDLA-extended *T. stictonotus* genome on the y-axis. Red lines indicate alignment in the forward direction, whereas blue lines indicate alignment in the reverse direction. LS: Linkage-group-extended (SELDLA-extended) Scaffold, chr: chromosome of FUGU5.

## Discussion

We initially speculated that whole-genome sequencing would be suitable to type SNPs in double haploid and/or haploid organisms because a single read would be sufficient. We note that SNPs are generally useful as polymorphic markers not only because they are easy to type, but also because they are very common. In addition, we can compensate for missing data at a site in one individual because the phase is successive except around crossover points, of which there were, on average, 1.3 per chromosome 1 in *T. rubripes* and 3.6 per chromosome 1 in *T. stictonotus.* In contrast, double haploid and/or haploid organisms are unsuitable for typing microsatellite polymorphic markers by conventional methods because they contain only half of the information compared with diploids. Moreover, we speculated that if we can trace the origin of each read in an organism to either of its parents, linkage analysis for the father and mother can be performed independently. This process is essentially equivalent to simultaneous construction of linkage maps for two lines of double haploid/haploid organisms. Therefore, it should also be possible to generate linkage maps even from low-coverage sequencing data. In support of this strategy, we also developed a new analytical pipeline, SELDLA, to generate high-resolution linkage maps, and, in parallel, to extend scaffolds/contigs based on thin, incomplete SNP data obtained from low-coverage genome data.

Following this approach, we mated two interfertile species, *T. rubripes* and *T. stictonotus,* to generate hybrid individuals with genes easily traceable to the parents. We succeeded in generating a new linkage map for *T. rubripes* from 442,723 SNPs in 188 hybrid individuals. These markers are significantly more than the number of crossover points in the sperm of the father, which was estimated to be several thousands. Therefore, the resolution of the linkage map was limited not by the number of markers, but by the number of crossover sites. Indeed, the number of SNP markers used in this analysis is the highest among any other combination of markers and typing methods. We also succeeded in generating a draft genome of higher quality than the current draft (FUGU5). For example, we mapped 95.7 % of scaffolds to the genome, of which 80 % were oriented, including those that were not previously located. In addition, we generated a linkage map for *T. stictonotus* for the first time, even with a very short N50 of 15 kb. In this case, we mapped 22.9 % of sequences to the genome.

Furthermore, we show in Fig. 4 that the genome we obtained can be extended, particularly at chromosome ends, which are problematic in FUGU5. Previously, the genetic length of chromosome 1 was reported as 99.7 cM and 179.9 cM for male and female *T. rubripes,* respectively(Kai et al. 2011), but because we mapped many more scaffolds near telomeres, we have extended the genetic length of chromosome 1 to 124.3 cM (Supplementary Fig. 2) and 353.4 cM (Supplementary Fig. 4), respectively. Of note, the reference sequence for SNP mapping was generated *de novo* by only one round of paired sequencing. However, combining a series of mate-pair and/or paired-end sequence data will generate a much longer N50, and enable precise and easy location of most longer scaffolds/contigs, as was achieved in *T. rubripes.*

PacBio and Oxford Nanopore sequencing have been recently demonstrated to generate much longer contiguous sequences. Several new techniques to locate sequences in the genome, including 10× Genomics and Irys, have also been established. These techniques also generate high-quality *de novo* genome sequences, and we anticipate that our strategy will be proved very useful when combined with these new methods. More importantly, our method is based on genetic mapping, whereas others are based on physical mapping. We note that regardless of advances in physical methods, a genetic map would still be required to compare with and evaluate physical maps, as well as for various types of genetic analysis.

To save cost and labor, several genotyping methods based on a small number of polymorphic sites have been developed, including RAD-seq and genotyping-by-sequencing. If an inbred line is obtained by subrearing, the inbred individuals are essentially clones of the same haplotype, which can then be easily determined. However, this approach is limited to model organisms such as mice and zebrafish that can be inbred. In addition, there seems to be no actual example of this approach, because it is time-consuming. Similarly, generating double haploids/haploids is often difficult, and may also limit applicability. On the contrary, our method is based on mating interfertile species, and therefore should be more widely applicable. In our experience, torafugu embryos before hatching already provide sufficient DNA for genotyping, suggesting that hatching is not even necessary, which would further broaden the mating possibilities.

Thus, our method is one of the most effective and promising approaches for obtaining genomes from organisms that produce many eggs or seeds in one generation, such as fish, mollusks, amphibians, insects, and plants. In addition, we may even be able to apply our method to viviparous organisms via *in vitro* fertilization and/or differentiation of induced pluripotent cells into germ cells. Accordingly, we are now applying or planning to apply our method to obtain genomes from various organisms.

## METHODS

### Fish and artificial insemination

Eggs from a female *T. stictonotus* caught on the day of the experiment were artificially inseminated with stored sperm obtained from a *T. rubripes* male in 2014. Hybrid fry were collected individually 10 days after fertilization (4~6 days after hatching). Because we lost the actual source of eggs, we caught another *T. stictonotus* female for genome sequencing.

### Library preparation and sequencing

Genomic DNA was individually extracted from 192 fry using DNeasy Blood and Tissue Kit (QIAGEN, Hilden, Germany). Libraries were prepared using Nextera DNA Sample Prep Kit (Illumina, San Diego, CA, USA) and Nextera Index Kit (Illumina). Libraries from 96 hybrid fry were then mixed at equal concentrations to obtain two pools (192 samples in total), which were processed and sequenced on a HiSeq 2500 Sequencer (Illumina). Four samples with few reads were excluded. After SNP calling, we excluded another two samples with many heterozygous SNPs because contamination was suspected.

### Genome assembly

*T. rubripes* and *T. stictonotus* adults were also sequenced to generate reference sequences with which to identify the source of each raw read in hybrid fry, and to call and map SNPs. Raw data were obtained from one round of sequencing on HiSeq 2500. For *T. rubripes,* the stored sperm was used as source of paternal genomic DNA. For *T. stictonotus,* a second individual as described above was used as the source of a mother-equivalent genome. Reads were assembled in CLC Assembly Cell 5.0.3 (QIAGEN), with an insert size of 100–800 bp and default values for all other parameters.

## SELDLA

We developed SELDLA, a novel data processing pipeline to construct a linkage map from genomic hybrids. SELDLA is available at http://www.suikou.fs.a.u-tokyo.ac.jp/software/SELDLA/, and is illustrated in Fig. 2. Briefly, we first combined the *T. rubripes* genome (FUGU4) and the *de novo T. stictonotus* genome assembled in this study. Reads from both adult fish and from 192 hybrid fry were then mapped onto the combined genome in BWA mem ver. 0.7.15, using “- t 4 - M” as the input parameter(Li and Durbin 2010). Uniquely mapped reads were extracted, sorted in samtools ver. 1.4(Li et al. 2009), and mined for SNPs in UnifiedGenotyper in GATK ver. 3.2–2(McKenna et al. 2010). SNPs were called with “-stand_emit_conf 0 - glm SNP”, even from just one read.

We excluded regions in the combined genome to which reads from both *T. stictonotus* and *T. rubripes* were mapped. We also excluded regions to which reads were mapped with more than four times the average depth. Each contig/scaffold was then phased, and linkage was measured by concordance rate of phases from all samples. In particular, the phase at two contigs/scaffolds was considered concordant when such contigs/scaffolds were completely linked, i.e., without crossover points, in 188 hybrids. In contrast, the concordance rate was close to 50 % for two contigs/scaffolds that were not linked. We note that satisfactory scaffold elongation was achieved by setting the concordance rate threshold to 90 % for male fish and to 70 % for female fish, as recombination rates are lower in male pufferfish than in female pufferfish(Kai et al. 2011).

### Comparative genome analysis

To generate Circos and dot plots, LAST ver. 861(Kielbasa et al. 2011) was used for homology search. LAST results were converted to tab-delimited text using suitable scripts, and regions with continuous alignment of at least 1,000 bp were obtained.

Circos plots were drawn in Circos ver. 0.69–5(Krzywinski et al. 2009) to compare the selected fragments against the 22 chromosomes in *T. rubripes* (FUGU5). All scaffolds were also compared in awk scripts, and dot plots were drawn in R based on LAST results.

### Linkage mapping

Physical distance was calculated directly by adding scaffold lengths. Linkage distance was calculated using phase information at both ends of the scaffold, with 1 cM representing recombination in 1 of 100 individuals. To exclude false recombination due to noise, we eliminated phases that did not match continuously for three or more scaffolds in each individual.

### Code availability

Code for SELDLA pipeline can be found at http://www.suikou.fs.a.u-tokyo.ac.jp/software/SELDLA/.

### Accession codes

WGS data are available from the DNA Data Bank of Japan (Accession number: DRA006412).

### Data availability

SELDLA-extended genomes can be found at http://www.suikou.fs.a.u-tokyo.ac.jp/software/SELDLA/data/.

## ACKNOWLEDGMENTS

Some computations were performed on the NIG supercomputer at the ROIS National Institute of Genetics.

## AUTHOR CONTRIBUTIONS

S.A. conceive and designed the entire studies. K.Y. conceived and designed the data processing strategy and developed concept of SELDLA. S.A. and K.Y. wrote the manuscript. S.W. supervised generation of hybrid *Takifugu.* M.M., S.K., K.S., and S.A. generated the hybrid individuals. Y.I., Y.S., S.M. and S.A. contributed sequencing and SNP typing. All authors agreed on the final version of the submitted paper.

---

**Supplementary Figure 1** Dot plot homology comparison of all chromosomes between SELDLA-extended FUGU4 and FUGU 5. Regions that could be aligned over 1 kb or more are plotted, with FUGU5 on the x-axis and SELDLA-extended FUGU4 on the y-axis. Red lines indicate alignment in the forward direction, whereas blue lines indicate alignment in the reverse direction.

**Supplementary Figure 2** Physical (Mb) and genetic (cM) distances in each SELDLA-extended *T. rubripes* chromosome. LS: Linkage-group-extended Scaffold corresponding to the same chromosome number of FUGU5.

**Supplementary Figure 3** Dot plot homology comparison of all chromosomes between FUGU 5 and SELDLA-extended *T. stictonotus* genome. Regions that could be aligned over 1 kb or more are plotted, with FUGU5 on the x-axis and SELDLA-extended *T. stictonotus* genome on the y-axis. Red lines indicate alignment in the forward direction, whereas blue lines indicate alignment in the reverse direction.

**Supplementary Figure 4** Physical (Mb) and genetic (cM) distances in each SELDLA-extended *T. stictonotus* chromosome. LS: Linkage-group-extended Scaffold corresponding to the same chromosome number of FUGU5.

